# Improved science communication and student gains from an undergraduate biomedical research experience

**DOI:** 10.1101/2023.10.17.561956

**Authors:** Donna Ward, Yueh-Ying Han, Christopher Qoyawayma, April Dukes, T. Brooke McClendon, Michelle L. Manni

**Author notes:** Corresponding Author: Michelle L. Manni. Authors contributed equally (Donna Ward, Yueh-Ying Han).

## Abstract

Summer research experiences expose undergraduate students to biomedical research in a laboratory or clinical setting, but often do not incorporate formal learning on scientific communication. Proper written and oral communication of science is essential to succeed in biomedical fields. This study examined whether participation in a science communication series (SCS) would increase gains in research and written communication abilities among students participated in the University of Pittsburgh School of Medicine Department of Pediatrics Summer Research Internship Program. Surveys were administered at the beginning and end of the program to evaluate their summer undergraduate research experience (SURE). Positive personal and professional gains in research and communication skills were identified through participation in both SURE and SCS. Participation in the SCS also significantly improved the quality and presentation of research abstracts. Focused learning in science communication during SUREs would improve undergraduate students’ personal and professional abilities in biomedical research and medicine.

## INTRODUCTION

Summer undergraduate research experiences (SUREs) are offered to increase student retention in science, technology, engineering, and mathematics (STEM) fields and enhance interest in careers in science and medicine (Bauer and Bennett 2003, Russell, Hancock et al. 2007, Tsui 2007). Although proper communication of one’s scientific or medical work is essential for careers in these fields, there is little focus on developing these skills while participating in SUREs. It is a common requirement at the conclusion of a SURE program for students to construct an abstract, poster, or oral presentation of their research; however, the necessary instruction on scientific communication is not always included in SURE curriculum. SUREs may be enhanced by incorporating curriculum that focuses on introducing and teaching science communication skills.

Most SURE programs target students in their sophomore to senior year of undergraduate studies. While it is universal for undergraduate students to receive instruction in English or Composition courses during their undergraduate training, it is less common for undergraduate science majors to take courses dedicated to technical or scientific writing (Jerde and Taper 2004, Fernández, García et al. 2018). Thus, these undergraduate students are likely to move on to advanced training in biomedical fields without fundamental knowledge of the tone, organization, and conciseness of scientific writing, all of which are prerequisites to professionally write papers, reports, or grant proposals. Instead of developing these skills in a classroom, they are more commonly acquired through trial and error during lab courses, a graduate program, or in their career. Without background and training in these areas, researchers are often hesitant to put themselves in situations requiring them to use scientific writing (Jerde and Taper 2004). Increasing undergraduate student exposure, especially within science majors, is critical to their post baccalaureate success.

Participation in SUREs is one way in which undergraduate students can gain essential research skills absent from their undergraduate curriculum. These skills include exposure to the practice of science in a research laboratory, being a member of a research team, research study design, data collection, and analysis and communication of scientific ideas and data (Weston and Laursen 2015). At the University of Pittsburgh School of Medicine Department of Pediatrics and Pittsburgh Children’s Hospital of Pittsburgh Summer Research Internship Program (SRIP), the goal of SRIP in the summer of 2021 was to create an immersive experience for undergraduate students in a research laboratory, allowing the students to gain knowledge and exposure to scientific thinking, methods, and techniques as well as emphasizing both written and verbal scientific communication. It is hypothesized that the incorporation of a scientific communication series (SCS) would result in enhanced written communication scores as well as student confidence in and satisfaction with their research experience and increase their desire to pursue a STEM advanced degree.

## METHODS

### Subjects

Fifty-seven undergraduate students, who were participating in SRIP, were asked to voluntarily participate in a survey at the start and end of their 8-week research experience. In 2021, students participating in this SURE were also provided the opportunity to participate in 6 optional sessions focused on learning about effective science communication. This study was reviewed and approved by the University of Pittsburgh Institutional Review Board (IRB# STUDY21040138).

### Survey

The following validated and published survey, ME-SALG: Student Assessment of their Learning Gains survey (McDevitt, Patel et al. 2016), was utilized to evaluate the experiences of the student participants in SRIP. Five sections of this questionnaire were included: Gains in thinking and working like scientist, Personal gains related to research work, Gains in skills, Attitudes or behaviors as a researcher, and Becoming a scientist (Wisconsin Center for Educational Research 2010). In addition, questions on the demographic information were included in the survey.

Data was collected twice during the SURE. Students completed the survey when they arrived prior to any learning sessions and at the conclusion of the program. In the post survey, students were asked if they participated in the SCS seminars or used any resources or materials provided via email to the students during their SURE.

### Science communication series (SCS) structure and participation

Undergraduate students participating in the SURE were given the opportunity to voluntarily participate in a six-session SCS during the SRIP. Each session was one hour in length, offered virtually using Microsoft Teams, and led by one or more facilitators of the SURE program. The seminar series focused on improving effective scientific communication through reading literature as well as communicating scientific ideas through writing and verbally. Seminar topics included: how to read and understand scientific literature, how to write a scientific abstract, how to design and present a scientific poster, how to provide constructive feedback on scientific writing, and how to orally present a summary of scientific research.

### Abstract Scoring Method

To assess if participation in the SCS would improve written communication in science, abstracts of students’ research projects were independently evaluated and scored by three faculty members in a biological science from different institutions. Each faculty member possessed a PhD and had experience composing and evaluating forms of scientific writing. Each abstract was assigned a unique ID linked to the corresponding survey described above for analysis purposes. Scorers were blinded to all identifying information, including author names, race, year of participation in the SURE, and participation status in the SCS. Each abstract was scored using a rubric which was modified from the Society for Academic Emergency Medicine Annual Meeting Scientific Abstract Scoring System (Society for Academic Emergency Medicine 2019). Abstracts were scored in 9 areas: Title, Introduction, Clarity of Objectives, Choice of Approach/Methods, Results, Conclusions, Interest and Importance of Topic, Overall Presentation and Length. Each area was awarded points from 0 to 3 as seen in the Supplemental Information. Total abstract scores ranged from 0 to 27 where higher scores represent a better quality of the abstract. Final abstract score was calculated by the mean of the scores from the 3 evaluators.

Abstracts from 2019 SURE participants were scored (n=49) using the abstract rubric (Supplemental Information), and this cohort of abstracts were written before the SCS was established. Additionally, abstracts from 2021 SURE participants (n=57) were scored using the same rubric. The 2021 abstract cohort includes sub-populations of abstracts from individuals who participated in SCS or used provided handouts and materials (n=34) and individuals who did not participate in SCS nor used SCS handouts and materials (n=13).

First, abstract scores were compared between those who participated in the 2019 SURE (n=49) and who participated in the 2021 SURE (n=57). Secondly, abstract scores were compared between students participated in SCS or used provided handouts and materials of 2021 SURE (n=34) and those students who participated in 2019 SURE (before SCS was established) and students who did not participate or use SCS materials in 2021 SURE (total n=72).

### Statistical Analyses

An overall score of each section from the Learning Gains survey was calculated by the mean of questions within the section. The Paired Samples t-Test (paired t-test) was used to compare the scores of pre- and post-survey taken from the same participant. Two-sample t-test was used for the comparison of the mean abstract scores between two groups. Two-sided *p*-values less than 0.05 were considered statistically significant. All analyses were conducted using Statistical Analysis System 9.4 software (SAS Institute, Inc, Cary, North Carolina).

## RESULTS

### Participants

The 2021 SURE Program hosted a total of 57 students, who were given the option to attend courses offered on science communication. Out of these 57 students, 30 students (52.6%) completed both the pre- and post-survey questionnaires. Among these 30 students, 25 students (83.3%) attended at least one SCS or used handouts from the seminar to complete their abstract (Table 1). Participants within the study in the summer of 2021 were mostly white, non-Hispanic, female, residents of the state of Pennsylvania, going into their junior or senior year of an undergraduate program, and with at least one parent having completed a post-graduate education (Table 1).

**Table 1.** Characteristics of participated students with pre- and post-survey.

| **Variable** | **All participants (n=30)** |
| --- | --- |
| Age | 20.7 ± 1.7 |
| Female gender | 21 (70.0) |
| Non-Hispanic | 28 (93.3) |
| Race |  |
| Asian | 7 (23.3) |
| Black | 1 (3.3) |
| White | 20 (66.7) |
| Other | 2 (6.7) |
| Year in College |  |
| Freshman | 3 (10.0) |
| Sophomore | 9 (30.0) |
| Junior | 9 (30.0) |
| Senior | 7 (23.3) |
| Other | 2 (6.7) |
| Highest parental education |  |
| High school/some college | 5 (16.7) |
| College | 7 (23.3) |
| Post-graduate | 18 (60.0) |
| Resident of Pennsylvania | 24 (80.0) |
| Participated in any section of science seminar | 25 (83.3) |
| Use any seminar materials |  |
| None | 5 (16.7) |
| 1 | 3 (10.0) |
| 2 | 8 (26.7) |
| 3 | 5 (16.7) |
| 4 | 5 (16.7) |
| 5 | 2 (6.7) |
| 6 | 2 (6.7) |

Results are shown as mean ± standard deviation (SD) for continuous variables, and as N (%) for categorical variables.

### Personal Gains

An overall significant increase (*p*<0.01) in personal gains related to research was observed between pre- and post-survey results among four sections (gains in thinking and working like scientist, personal gains related to research work, gains in skills, and attitudes or behaviors as a researcher, Figure 1 and Table 2). Comparison between pre- and post-survey for individual questions in each section are also reported (Supplemental Table 1-5). While an increase in gains in all areas was observed, a significant increase was noted in the following categories: confidence in ability to contribute to science, comfort in discussing scientific concepts with others, developing patience with the slow pace of research, understanding what research work is like and taking greater care in conducting procedures in the lab or field.

**Table 2.** Summary of all the overall scores (n=30).

| **Overall score for each section** | **Pre-survey** | **Post-survey** | **P-value\*** |
| --- | --- | --- | --- |
| Gains in thinking and working like scientist | **3.63 ± 0.65** | **4.52 ± 0.46** | **<0.01** |
| Personal gains related to research work | **3.75 ± 0.65** | **4.53 ± 0.58** | **<0.01** |
| Gains in skills | **3.46 ± 0.69** | **4.12 ± 0.72** | **<0.01** |
| Attitudes or behaviors as a researcher | **3.21 ± 0.96** | **4.30 ± 0.66** | **<0.01** |
| Becoming a scientist | 2.80 ± 0.54 | 2.81 ± 0.52 | 0.87 |

Results are shown as mean ± standard deviation (SD).
\* Comparison between pre-survey and post-survey was analyzed by paired t-test.

**Figure 1.**
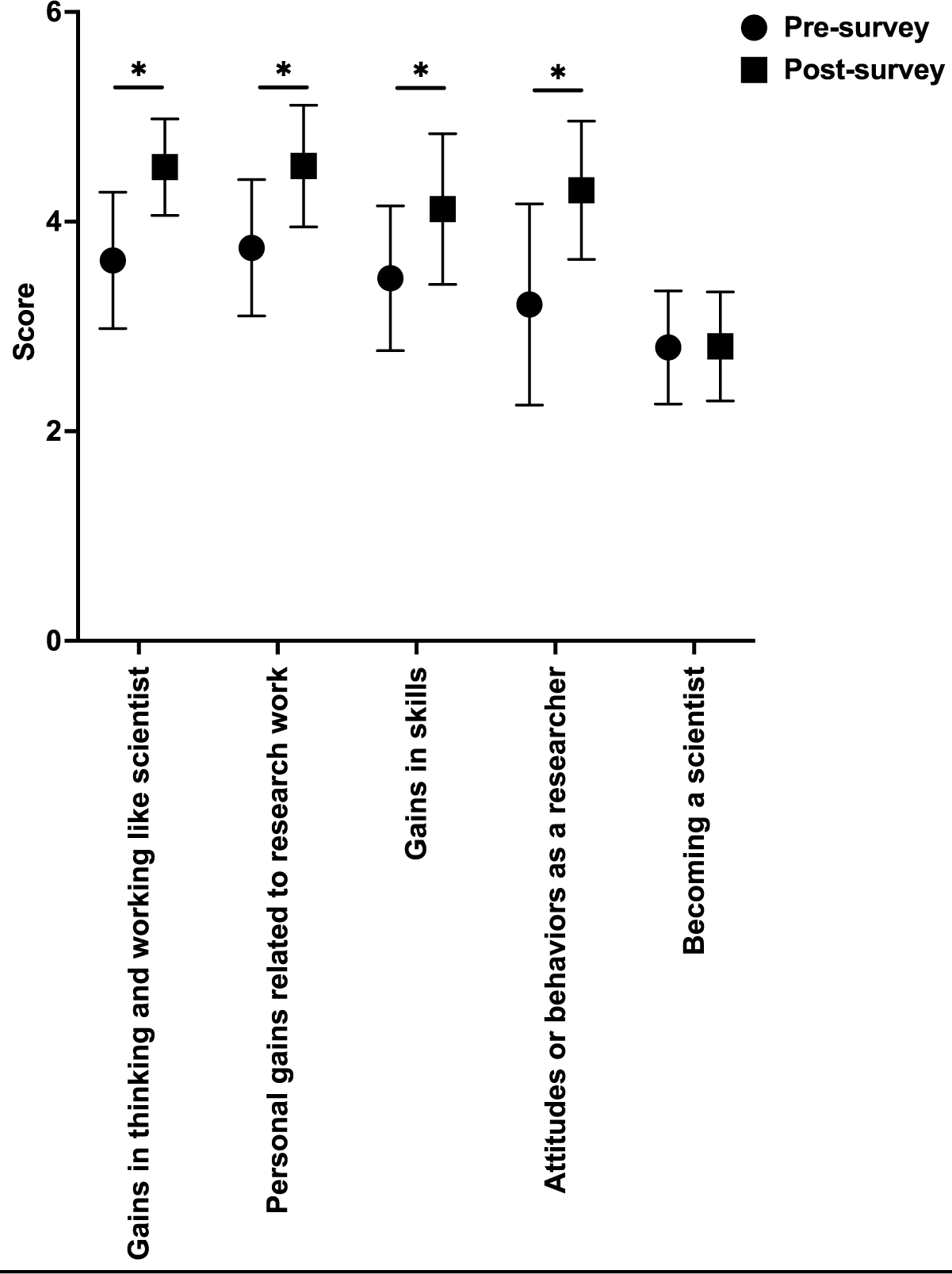
Mean overall changes in learning gains between pre- and post-SURE. Data shown is mean +/- standard deviation. \**p*<0.01, paired t-test

Significant gains were also observed in writing scientific reports or papers, defending an argument when asked questions, explaining my project to people outside of my field, preparing a scientific poster, conducting observations in the lab or field and understanding journal articles. A significant gain overall and in each item relating to attitudes or behaviors as a researcher was also observed, however, no significant gains were noted between pre- and post-survey results, indicating student participants had clear career aspirations and the SURE did not alter their desire to become a scientist.

These results were also validated by survey responses from 2022, where comparable results were found even with a small sample size and more diverse population of undergraduate student participants (Supplemental Table 6).

### Abstract Scores

Although not statistically significant, the average total abstract score was slightly higher among students participating in the 2021 SURE program as compared to those who participated in the 2019 SURE program (20.3 ± 3.1 vs. 19.0 ± 3.6, P=0.07, Table 3). Students who participated in the SCS or used provided handouts and materials from 2021 SURE had a significantly higher average abstract score as compared to those did not (20.5 ± 2.8 vs. 19.1 ± 3.5, P=0.04, Table 4). Specifically, scores for introduction, objective, and presentation were significantly higher in students participating in the 2021 SURE program and in those who participated in SCS or used related materials (Table 3 and Table 4).

**Table 3.** Comparison between abstract scores by the year of participation.

| Average score | Students from 2019 | Students from 2021 | P-value* |
| --- | --- | --- | --- |
|  | (n=49) | (n=57) |  |
| **Total scores** | **19.0 ± 3.6** | **20.1 ± 3.1** | **0.07** |
| Title | 2.5 ± 0.4 | 2.4 ± 0.5 | 0.17 |
| Introduction | 2.2 ± 0.9 | 2.5 ± 0.6 | 0.06 |
| Objective | 2.2 ± 0.7 | 2.5 ± 0.5 | **0.03** |
| Methods | 2.3 ± 0.7 | 2.3 ± 0.7 | 0.94 |
| Results | 2.2 ± 0.9 | 2.2 ± 0.9 | 0.82 |
| Conclusion | 1.8 ± 1.0 | 1.8 ± 0.9 | 0.95 |
| Importance | 1.8 ± 0.4 | 1.9 ± 0.4 | 0.27 |
| Presentation | 2.3 ± 0.5 | 2.5 ± 0.4 | **0.006** |
| Length | 1.5 ± 1.1 | 1.9 ± 1.0 | 0.06 |

Results are shown as mean ± standard deviation (SD).
\* t-test was used for the comparison between two groups

**Table 4.** Comparison of abstract scores by participation in SCS or use of SCS materials.

| Abstract score | Students from 2019 or<br>those from 2021 that<br>did not participate in<br>SCS nor used<br>materials (n=72) | Student from 2021<br>that participated SCS<br>or used materials<br>(n=34) | P-value* |
| --- | --- | --- | --- |
| **Total scores** | **19.1 ± 3.5** | **20.5 ± 2.8** | **0.04** |
| Title | 2.5 ± 0.5 | 2.4 ± 0.5 | 0.13 |
| Introduction | 2.3 ± 0.9 | 2.6 ± 0.4 | **0.03** |
| Objective | 2.3 ± 0.6 | 2.6 ± 0.5 | **0.03** |
| Methods | 2.2 ± 0.7 | 2.3 ± 0.7 | 0.51 |
| Results | 2.2 ± 0.9 | 2.3 ± 0.9 | 0.57 |
| Conclusion | 1.8 ± 1.0 | 2.0 ± 0.7 | 0.12 |
| Importance | 1.8 ± 0.4 | 2.0 ± 0.4 | 0.09 |
| Presentation | 2.3 ± 0.5 | 2.6 ± 0.4 | **0.008** |
| Length | 1.6 ± 1.1 | 1.8 ± 1.0 | 0.35 |

Results are shown as mean ± standard deviation (SD).
\* t-test was used for the comparison between two groups

## DISCUSSION

The SCS and SURE had a positive impact on student’s self-reported professional and personal gains in research skills and confidence in their scientific abilities. The SCS provided statistically significant gains in student abilities in scientific communication. Even though the SURE did not impact students’ desired career path, this assessment provided strong evidence that incorporation of SCS into SUREs would be beneficial for student participants’ learning and may enhance their abilities moving into a biomedical career.

The study included students in all years of their undergraduate studies, with many students being rising juniors and seniors in the Fall of 2021. Due to the number of upperclassman students involved in this study, as well as the higher education level of student family members, and the main career goal of this student population being to attend medical school, there is an increased possibility that some of these students had previous research experience. As upperclassmen, these students are more likely to have taken more scientific and laboratory courses during their undergraduate training, that may have increased their baseline knowledge and comfort levels within a research setting. Further, many students were already interested in STEM research and careers and stated that they were focused on pursuing medical school education in the future. Although previous research experience and career objectives may have inflated the personal and professional gains in the pre-surveys, there were still robust, significant gains in professional and personal skills obtained by participating in the 2021 SRIP (Tables 2 and 4). Thus, additional summer research experience is still beneficial for gaining further confidence and skills to pursue and retain students in biomedical fields.

To assess the effectiveness of the SCS, abstracts submitted in 2019 and 2021 were scored by three independent PhD-trained scientists. These results revealed that formal instruction of science communication will improve written communication of one’s science during SUREs but also increased student gains overall in the SURE. While a structured rubric for scoring abstracts was used by the reviewers, there may be some subjectiveness to assigning scores. Future study incorporates training for scoring and assessing agreement between evaluators will help to prevent potential bias.

Lastly, the number and diversity of student participants was limited in our cohort (Table 1). Although it is reported that underrepresented minorities tend to change to non-STEM related majors during their undergraduate career and only represent 11% of those in STEM related careers (Tsui 2007), a focus needs to be placed on recruiting more students of diverse backgrounds into SUREs. As previous studies have shown that participation in undergraduate research programs increases student enrollment and completion of graduate programs (Bauer and Bennett 2003), methods to increase student diversity in SUREs should be explored and should be a focus to increase diversity in STEM fields. Our study would suggest that students who participate in SUREs not only gain exposure and experience in a research lab, but also valuable professional and personal skills for their future in a biomedical career. Collecting and incorporating data from all summer undergraduate research programs offered across the University of Pittsburgh in a variety of scientific disciplines may be beneficial for future analyses.

## REFERENCES

Bauer, K. and J. Bennett (2003). “Alumni perceptions used to assess undergraduate research experience.” The Journal of Higher Education 74(2): 210–230.

Fernández, E., et al. (2018). “Students’ satisfaction and perceived impact on knowledge, attitudes and skills after a 2-day course in scientific writing: a prospective longitudinal study in Spain.” BMJ Open 8(1): e018657.

Jerde, C. and M. Taper (2004). “Preparing Undergraduates for Professional Writing: Evidence Supporting the Benefits of Scientific Writing within the Biology Curriculum.” Journal of College Science Teaching 33(7): 34–37.

McDevitt, A. L., et al. (2016). “Insights into Student Gains from Undergraduate Research Using Pre- and Post-Assessments.” Bioscience 66(12): 1070–1078.

Russell, S., et al. (2007). “Benefits of undergraduate research experiences.” Science (Washington) 316(5824): 548–549.

Society for Academic Emergency Medicine (2019). “Society for Academic Emergency Medicine Annual Meeting Scientific Abstract Scoring System.” from https://www.saem.org/docs/default-source/annual-meeting/2019-annual-meeting/education/abstract-scoring-8-22-18.pdf?sfvrsn=ad3b09fd_2.

Tsui, L. (2007). “Effective strategies to increase diversity in STEM fields: a review of the research literature.” The Journal of Negro Education: 555-581.

Weston, T. and S. Laursen (2015). “The Undergraduate Research Student Self-Assessment (URSSA): Validation for Use in Program Evaluation.” CBE-Life Sciences Education 14: 1–10.

Wisconsin Center for Educational Research (2010). “SALG - Student Assessment of their Learning Gains: PREVIEW INSTRUMENT URSSA MASTER - REVIEW COPY, ORIGINAL.” Retrieved May 14, 2020, from https://www.colorado.edu/eer/sites/default/files/attached-files/urssa_master_reviewcopy.pdf

