## Supplemental Information for "Improved science communication and student gains from an undergraduate biomedical research experience"

**Abstract Scoring Method**

Method was modified from SAEM 2019 Annual Meeting scientific abstract scoring system) (https://www.saem.org/docs/default-source/annual-meeting/2019-annual-meeting/education/abstract-scoring-8-22-18.pdf?sfvrsn=ad3b09fd_2)

**TITLE** - Is the title specific, adequate and concise? Does it accurately describe the population studied, the study design or method of data collection or analysis, the research objective or question?

| | |
|---|---|
| 3 | The title perfectly describes the study, population, methods, and research question or objective. |
| 2 | The title describes the study very well. The title may be missing one of the following necessary elements: the population, methods, research question, or objective. |
| 1 | The title generally describes the study but may not mention more than one of the following: populations, main methods, research question or objective. |
| 0 | The title is vague and did not adequately describe the study presented in the abstract. |

**INTRODUCTION -** Is the context or need of the study made clear? Is the scientific rationale clearly stated?

| | |
|---|---|
| 3 | The relevant background and context of the study are clearly presented. |
| 2 | Most of the relevant background and context of the study are presented well but may be missing background on one key element. Scientific rationale for the project is clear. |

| | |
|---|---|
| 1 | Background or context of the study are stated. Many areas of the background may be sparse or unconnected. The scientific rationale is unclear. |
| 0 | The background and rationale for the work are largely absent or poorly written. |

**CLARITY OF OBJECTIVES -** Does the study have clear objectives (whether descriptive or hypothesis-testing)?

| | |
|---|---|
| 3 | Objective(s) of the study are well thought out and clearly stated. Hypothesis-driven objectives are testable. |
| 2 | Study objectives are stated. Objective could use some fine-tuning for clarity or to improve testability. |
| 1 | Stated objectives were poorly chosen, or stated hypothesis was difficult to test. |
| 0 | No clear objectives or hypothesis. |

**CHOICE OF APPROACH/METHODS** - Does this study use the right research methods for the scientific question? Are the Methods clearly described? Are the data sources clearly specified? Are the methods, analytical techniques and software tools specified where applicable for the project? Are the methods appropriate to the question being investigated?

| | |
|---|---|
| 3 | Chosen study design was the best feasible method for testing the stated hypothesis/ objectives (i.e., a robust design). |
| 2 | Chosen study design was sub-optimal but did test the stated hypothesis/objectives (i.e., an acceptable design). |

| | |
|---|---|
| 1 | Study design was unclear from the abstract, or it was unclear how the design tested the stated hypothesis/objectives. |
| 0 | Design did not test stated hypothesis/objectives. |

**RESULTS -** Are results available and described appropriately? **Abstracts should not say only that 'results will be presented'.

| | |
|---|---|
| 3 | The results are clearly stated and are perfectly described with very clear expression of ideas. |
| 2 | The results are generally well-written. Not all results may be discussed, or the results contains one or two confusing or absent statements or concepts. |
| 1 | The results are poorly written and leaves room for confusion many statements or concepts. |
| 0 | The results are largely (or fully) absent. It is unclear what was accomplished for this project. |

**CONCLUSIONS** - Are the conclusions clear and concise? Do they reflect the aims and objectives? Are they supported by the results presented? Where applicable, are key study limitations acknowledged? Where appropriate, are the implications made clear for policy, practice and further research?

| | |
|---|---|
| 3 | The conclusions are clearly stated, full support results presented, and acknowledge limitations and future directions. |
| 2 | The conclusions are stated, but could use further clarification, elaboration, or brevity. Authors mention either limitations or future directions. |

| | |
|---|---|
| 1 | The conclusions are mentioned but do not fully support results presented. Authors mention neither limitations nor future directions. |
| 0 | Conclusions are absent, or it is largely unclear what the authors conclude. |

**INTEREST and IMPORTANCE OF TOPIC -** Is it interesting? Does it have the potential to create impact (i.e. change clinical or public health practice or policy, improve health, reduce inequalities in health, change the course of science)? Is it novel/exciting/much better methodologically than other studies in the area?

| | |
|---|---|
| 3 | This topic, or its foreseeable progeny, is relevant to physicians or scientists, or is highly innovative. |
| 2 | This is an important topic that will lead to information of interest to many or most physicians or scientists, including those who do not study this topic. |
| 1 | This is a topic that will be interesting to some physicians or scientists but may mostly impact those who study this topic. |
| 0 | This topic is only of interest to the small group of people who study it and is unlikely to result in important knowledge. |

**OVERALL PRESENTATION** - Does this abstract reflect high-quality writing and attention to detail?

| | |
|---|---|
| 3 | Perfect grammar, no errors, very clear expression of ideas. |
| 2 | Generally well-written, but leaves room for confusion on some concepts or has one or two errors. |

| | |
|---|---|
| 1 | Fairly written, but contains noticeable grammar and/or spelling errors. Abstract is at times confusing or hard to follow. |
| 0 | Poorly written. Hard to understand, idiosyncratic phrasing, or awkward abbreviations. |

**LENGTH OF ABSTRACT** - Is the abstract an appropriate length (approximately 250-350 words)?

| | |
|---|---|
| 3 | Abstract length is appropriate (250-350 words). Fits on one page and has an excellent balance of information and succinctness. |
| 2 | Generally sufficient abstract length but may too short/missing information or too long/repetitive. Abstract word count is between 150-250 or 350-400 and fits to one page. |
| 1 | The abstract length may be significantly shorter than expected with missing information or slightly over with repetitive phrases and/or statements. Abstract word count is 100-150 or 400-500 and fits to one page. |
| 0 | Abstract is over 500 words, under 100 words, or over a page in length. |
