## Supplemental Table 6 for "Improved science communication and student gains from an undergraduate biomedical research experience"

**Supplemental Table 6- Undergraduate research student self-assessment results for 2022 (n=15)**

| **Items** | **Pre-survey** | **Post-survey** | **P-value***|
|---|---|---|---|
| Gains in thinking and working like scientist | 3.38 ± 0.90 | 4.38 ± 0.59 | **<0.01** |
| Personal gains related to research work | 3.57 ± 0.86 | 4.40 ± 0.60 | **<0.01** |
| Gains in skills | 3.43 ± 0.62 | 4.12 ± 0.57 | **0.02** |
| Attitudes or behaviors as a researcher | 2.95 ± 0.88 | 4.40 ± 0.51 | **<0.01** |
| Becoming a scientist | 2.83 ± 0.63 | 2.84 ± 0.61 | 0.82 |

Results are shown as mean ± standard deviation (SD).

* Comparison between pre-survey and post-survey by paired t-test.
