## Supplemental Table 5 for "Improved science communication and student gains from an undergraduate biomedical research experience"

**Supplemental Table 5- Becoming a scientist (n=30)**

| Items | Pre-survey | Post-survey | P-value |
|---|---|---|---|
| 1. Enroll in a Ph.D. program in science, mathematics, or engineering? | 2.87 ± 1.38 | 2.97 ± 1.40 | 0.56 |
| 2. Enroll in a master program in science, mathematics, or engineering? | 2.60 ± 1.35 | 2.70 ± 1.39 | 0.66 |
| 3. Enroll in a combined M.D/Ph.D program | 2.57 ± 1.19 | 2.67 ± 1.35 | 0.61 |
| 4. Enroll in medical or dental school? | 4.10 ± 1.49 | 4.07 ± 1.51 | 0.57 |
| 5. Enroll in a program to earn a different professional degree (i.e. law, veterinary medicine, etc.) | 1.90 ± 1.40 | 1.67 ± 1.03 | 0.25 |
| 6. Pursue certification as a teacher? | 1.93 ± 1.11 | 1.97 ± 1.33 | 0.81 |
| 7. Work in a science lab? | 3.57 ± 1.01 | 3.67 ± 1.22 | 0.63 |
| **Overall becoming a scientist** | **2.80 ± 0.54** | **2.81 ± 0.52** | **0.87** |

Results are shown as mean ± standard deviation (SD).

* Comparison between pre-survey and post-survey by paired t-test.
