## Supplemental Table 4 for "Improved science communication and student gains from an undergraduate biomedical research experience"

**Supplemental Table 4- Attitudes or behaviors as a researcher (n=30)**

| Items | Pre-survey | Post-survey | P-value* |
|---|---|---|---|
| 1. Engage in real-world science research | 3.40 ± 1.25 | 4.63 ± 0.61 | <0.01 |
| 2. Feel like a scientist | 3.20 ± 1.24 | 4.47 ± 0.82 | <0.01 |
| 3. Think creatively about the project | 3.57 ± 1.07 | 4.23 ± 0.77 | <0.01 |
| 4. Try out new ideas or procedures on your own | 2.60 ± 1.22 | 3.63 ± 1.13 | <0.01 |
| 5. Feel responsible for the project | 3.20 ± 1.24 | 4.33 ± 0.88 | <0.01 |
| 6. Feel a part of a scientific community | 3.27 ± 1.26 | 4.50 ± 0.86 | <0.01 |
| **Overall attitudes or behaviors as a researcher** | **3.21 ± 0.96** | **4.30 ± 0.66** | **<0.01** |

Results are shown as mean ± standard deviation (SD).

* Comparison between pre-survey and post-survey by paired t-test.
