## Supplemental Table 3 for "Improved science communication and student gains from an undergraduate biomedical research experience"

**Supplemental Table 3- Gains in skills (n=30)**

| Items | Pre-survey | Post-survey | P-value* |
| --- | --- | --- | --- |
| 1. Writing scientific reports or papers | 3.07 ± 1.11 | 4.00 ± 1.11 | <0.01 |
| 2. Making oral presentations | 3.67 ± 0.92 | 4.00 ± 1.13 | 0.12 |
| 3. Defending an argument when asked questions | 3.43 ± 0.94 | 4.00 ± 0.87 | <0.01 |
| 4. Explaining my project to people outside my field | 3.28 ± 1.13 | 4.47 ± 0.57 | <0.01 |
| 5. Preparing a scientific poster | 2.97 ± 1.10 | 4.50 ± 0.97 | <0.01 |
| 6. Keeping a detailed lab notebook | 3.47 ± 1.14 | 4.15 ± 1.06 | 0.04 |
| 7. Conducting observations in the lab or field | 3.53 ± 0.94 | 4.25 ± 0.75 | <0.01 |
| 8. Using statistics to analyze data | 3.30 ± 1.12 | 3.93 ± 1.36 | 0.03 |
| 9. Calibrating instruments needed for measurement | 3.10 ± 1.06 | 3.34 ± 1.40 | 0.35 |
| 10. Working with computers | 3.80 ± 1.06 | 3.93 ± 1.11 | 0.63 |
| 11. Understanding journal articles | 3.73 ± 0.94 | 4.37 ± 0.67 | <0.01 |
| 12. Conducting database or internet searches | 3.73 ± 0.98 | 4.27 ± 0.94 | 0.02 |
| 13. Managing my time | 3.93 ± 0.83 | 4.28 ± 0.92 | 0.15 |
| **Overall gains in skills** | **3.46 ± 0.69** | **4.12 ± 0.72** | **<0.01** |

Results are shown as mean ± standard deviation (SD).

* Comparison between pre-survey and post-survey by paired t-test.
