## Supplemental Table 2 for "Improved science communication and student gains from an undergraduate biomedical research experience"

**Supplemental Table 2- Personal gains related to research work (n=30)**

| Items | Pre-survey | Post-survey | P-value* |
|---|---|---|---|
| 1. Confidence in my ability to contribute to science | 3.53 ± 0.94 | 4.40 ± 0.93 | <0.01 |
| 2. Comfort in discussing scientific concepts with other | 3.77 ± 0.86 | 4.37 ± 0.81 | <0.01 |
| 3. Comfort in working collaboratively with others | 4.27 ± 0.69 | 4.50 ± 0.73 | 0.18 |
| 4. Confidence in my ability to do well in future science courses | 4.07 ± 0.87 | 4.48 ± 0.91 | 0.12 |
| 5. Ability to work independently | 4.20 ± 0.71 | 4.38 ± 0.94 | 0.39 |
| 6. Developing patience with the slow pace of research | 3.40 ± 1.13 | 4.43 ± 0.82 | <0.01 |
| 7. Understanding what everyday research work is like | 3.13 ± 1.28 | 4.90 ± 0.40 | <0.01 |
| 8. Taking greater care in conducting procedures in the lab or field | 3.60 ± 0.97 | 4.77 ± 0.50 | <0.01 |
| **Overall personal gains related to research work** | **3.75 ± 0.65** | **4.53 ± 0.58** | **<0.01** |

Results are shown as mean ± standard deviation (SD).

* Comparison between pre-survey and post-survey by paired t-test.
