## Supplemental Table 1 for "Improved science communication and student gains from an undergraduate biomedical research experience"

SUPPLEMENTAL TABLES:

**Supplemental Table 1- Gains in thinking and working like scientist (n=30)**

| Items | Pre-survey | Post-survey | P-value* |
|---|---|---|---|
| 1. Analyzing data for patterns | 3.67 ± 0.92 | 4.40 ± 0.89 | <0.01 |
| 2. Figuring out the next step in a research project | 3.03 ± 0.96 | 4.60 ± 0.72 | <0.01 |
| 3. Problem-solving in general | 4.30 ± 0.84 | 4.57 ± 0.73 | 0.10 |
| 4. Formulating a research question that could be answered with data | 3.53 ± 0.94 | 4.37 ± 0.81 | <0.01 |
| 5. Identifying limitations of research methods and designs | 3.37 ± 0.93 | 4.43 ± 0.68 | <0.01 |
| 6. Understanding the theory and concepts guiding my research project | 3.72 ± 1.00 | 4.76 ± 0.44 | <0.01 |
| 7. Understanding the connections among scientific disciplines | 3.63 ± 0.85 | 4.43 ± 0.68 | <0.01 |
| 8. Understanding the relevance of research to my coursework | 3.80 ± 0.96 | 4.62 ± 0.68 | <0.01 |
| **Overall gains in thinking and working like scientist** | **3.63 ± 0.65** | **4.52 ± 0.46** | **<0.01** |

Results are shown as mean ± standard deviation (SD).

* Comparison between pre-survey and post-survey by paired t-test.
